# Lectin-Assisted Imaging Mass Cytometry (Lectin-IMC) Enables Spatial Validation of Disease-Associated Glyco-Niches within a Multimodal Glycoprotein Analysis Framework

**DOI:** 10.64898/2026.09.19.752828

**Authors:** Chiaki Nagai-Okatani, Umeko Horiuchi, Patcharaporn Boottanun, Kei Miyako, Atsushi Kuno

**Author notes:** Correspondence (C.N-O.).

## Abstract

Understanding the pathological significance of protein glycosylation and translating disease-associated glycan alterations into diagnostic and therapeutic opportunities require integrated analysis of glycans, their carrier glycoproteins, and spatial context. Here, we present an expanded multimodal glycoprotein analysis framework incorporating lectin-assisted imaging mass cytometry (Lectin-IMC) for stepwise discovery and spatial validation of disease-associated glyco-niches. Disease-associated glycans identified by laser microdissection–assisted lectin microarray (LMD-LMA) tissue glycome mapping are spatially evaluated by Lectin-IMC in relation to cell types and tissue microenvironments, and candidate carrier glycoproteins identified by MS-based glycoproteomics are subsequently incorporated for higher-order evaluation of glycan–protein–cell type relationships. Multiplex panel design is guided by LMA-based assessment of lectin–lectin interactions and biologically informed probe selection. Using a dilated cardiomyopathy model, we constructed a five-lectin panel centered on Wisteria floribunda agglutinin (WFA), which recognizes fibrosis-associated asialo N-glycans identified previously. Glycan-dependent WFA detection was validated by competitive inhibition and PNGase F treatment, and WFA-reactive glycans were spatially associated with fibrotic regions containing ACTA2⁺VIM⁺ myofibroblast-like cells. Among six extracellular matrix glycoprotein candidates identified in WFA-binding fractions, periostin showed the strongest spatial correspondence with WFA-positive regions by pixel-based quantitative analysis and was further associated with ACTA2⁺VIM⁺WFA⁺ fibrotic microenvironments. The optimized Lectin-IMC panel was also transferable to a human FFPE cardiomyopathy specimen. Collectively, Lectin-IMC provides intuitive spatial visualization and interpretability of glycan–protein–cell type relationships and serves as an on-tissue spatial validation and prioritization layer within an iterative multimodal framework in which spatially defined glyco-niches can guide subsequent proteomic or glycoproteomic discovery.

## INTRODUCTION

Protein glycosylation plays critical roles in regulating protein structure and function and in mediating cell–cell and cell–matrix interactions.^1^ Disease-associated alterations in glycan structures have therefore attracted increasing attention as potential diagnostic markers and therapeutic targets.^2,3^ Recent advances in glycomic and glycoproteomic technologies, including mass spectrometry (MS) and lectin microarrays (LMAs), have enabled increasingly comprehensive profiling of disease-associated glycan alterations and their carrier glycoproteins, accelerating glycan-based biomarker discovery and glycan-targeted therapeutic research.^4–6^ However, understanding the biological and pathological significance of glycan alterations requires not only characterization of glycan structures themselves but also identification of the glycoproteins that carry these glycans and determination of how these molecular features are organized within tissue microenvironments.^7,8^

These requirements are particularly important for defining disease-associated “glyco-niches,” in which glycan features, their candidate carrier glycoproteins, and surrounding cellular and extracellular components are spatially integrated.^8^ A glyco-niche cannot be defined by glycan localization alone because the same glycan motif can be carried by multiple glycoproteins and may be distributed across distinct cellular and extracellular compartments. Characterization of such niches therefore requires coordinated information on (i) disease-associated glycan features, (ii) candidate carrier glycoproteins associated with those features, and (iii) their cellular and spatial context within tissues. These information layers have traditionally been acquired using different analytical platforms, and practical approaches that reconnect them within the same pathological context remain limited.^6–8^

In parallel, rapid developments in single-cell and spatial omics, including spatial transcriptomics and MS-based spatial proteomics, have substantially improved our understanding of cell-type-specific molecular organization in tissues.^9,10^ Nevertheless, most spatial omics approaches primarily rely on gene or protein expression measurements and generally lack direct information on glycan structures. Recent advances have begun to address this gap through complementary strategies, including highly multiplexed lectin-based imaging,^11^ near-cellular-resolution *N*-glycan mass spectrometry imaging,^1^^2^ integration of glycan information with multiplex protein imaging or spatial multiomics,^13,14^ and spatially resolved glycoproteomics guided by *N*-glycan imaging.^15^ These studies demonstrate that glycan information can be incorporated into spatial analysis together with protein or cellular information. However, an analytical framework that begins with upstream discovery of disease-associated glycans and candidate carrier glycoproteins and then reconnects these candidates through multiplex on-tissue validation of glycan–protein–cellular relationships remains underdeveloped. Thus, the analytical gap is not simply the ability to perform multiplex glycan imaging, but the ability to integrate discovery and spatial validation across glycan, glycoprotein, and tissue-context information.

This challenge is particularly relevant in extracellular microenvironments such as fibrotic tissues, where pathological molecular events are not confined to individual cells but emerge as spatially organized structures composed of secreted and accumulated extracellular matrix (ECM) molecules.^16^ Fibrosis represents a typical example in which ECM proteins produced by activated stromal cell populations, such as myofibroblasts, accumulate within tissues and form characteristic pathological microenvironments.^17^ In these settings, disease-associated glycan alterations may be linked to specific ECM glycoproteins and to the cellular populations responsible for their production or deposition. We recently established a hierarchical lectin-based spatial glycomics workflow combining laser microdissection–assisted lectin microarray (LMD-LMA) tissue glycome mapping, lectin histochemistry, and low-vacuum scanning electron microscopy (LVSEM), which enabled ultrastructural localization of fibrosis-associated glycan signals within ECM-rich cardiac tissue.^18^ This workflow resolved the structural context of disease-associated glycans but did not directly connect these signals to specific cellular populations or prioritize candidate carrier glycoproteins within the same tissue context.

In this study, we therefore extend this multimodal glycoprotein analysis framework by incorporating lectin-assisted imaging mass cytometry (Lectin-IMC) for stepwise discovery and characterization of disease-associated glyco-niches (Figure 1A). The framework integrates LMD-LMA tissue glycome mapping for discovery of disease-associated glycan alterations,^19,20^ MS-based glycoproteomics for identification of candidate carrier glycoproteins,^6,7^ and Lectin-IMC for multiplex evaluation of glycan–protein–cell type relationships in situ. Lectin-IMC panel design is further supported by LMA-based assessment of lectin–lectin interactions and by selection of lectin probes, candidate carrier protein markers, and cell-type markers according to the biological and pathological information required to interpret the target microenvironment. Using a dilated cardiomyopathy model, we applied this framework to fibrosis-associated asialo N-glycans recognized by Wisteria floribunda agglutinin (WFA) and to candidate ECM glycoproteins previously identified in WFA-binding fractions by MS-based glycoproteomic analysis.^21^ We further evaluated the transferability of the optimized Lectin-IMC panel to a human formalin-fixed paraffin-embedded (FFPE) cardiomyopathy specimen. By positioning Lectin-IMC not as an isolated multiplex glycan-imaging method but as an on-tissue validation and prioritization layer within a broader discovery–validation workflow, this framework provides a practical strategy for spatially prioritizing disease-associated glycoproteins and may provide spatial entry points for subsequent image-guided proteomic or glycoproteomic analysis.

**Figure 1.**
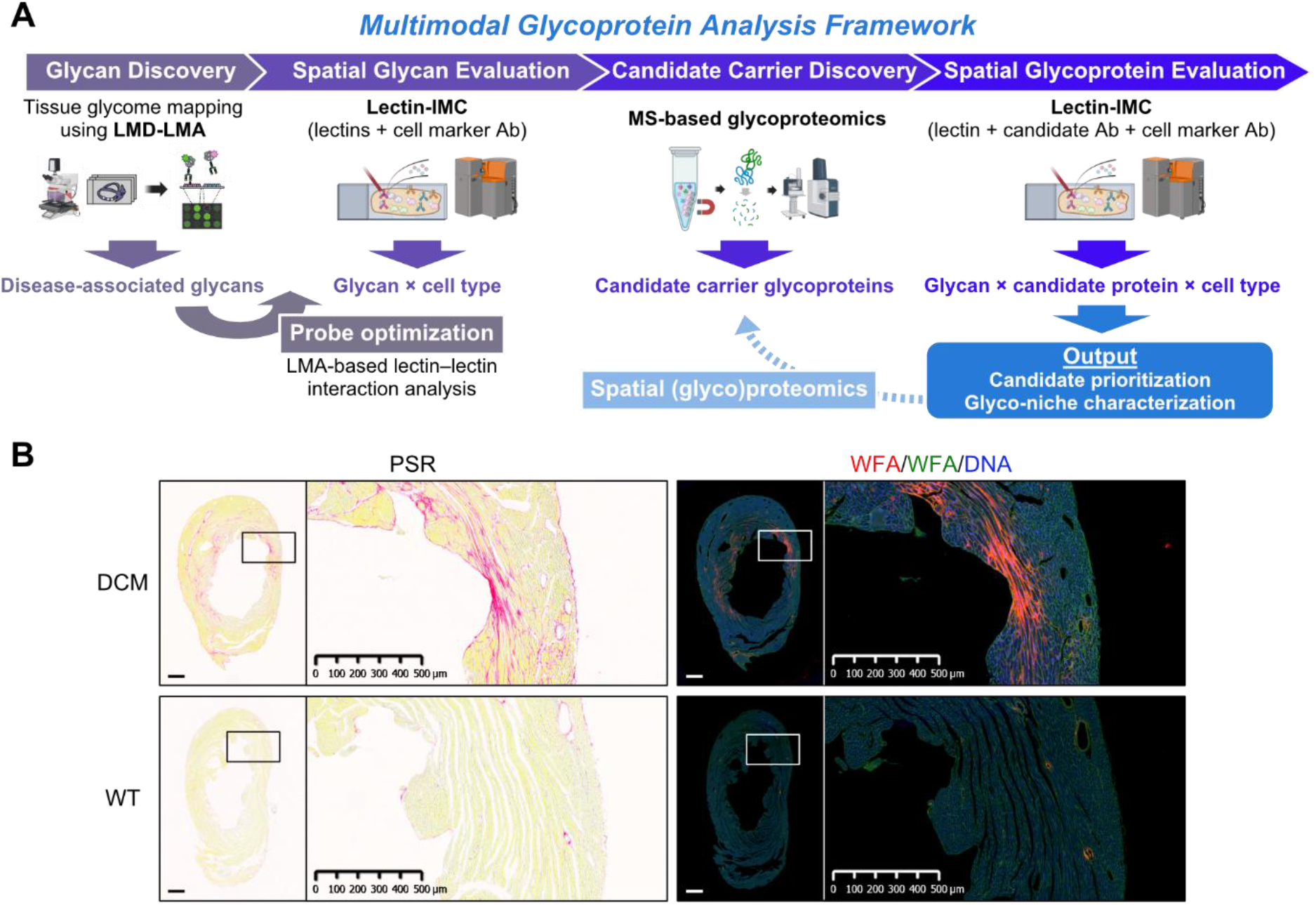
Multimodal glycoprotein analysis framework for spatial validation of disease-associated glyco-niches. (A) Schematic overview of the multimodal glycoprotein analysis framework. Disease-associated glycans identified by LMD-LMA tissue glycome mapping are first spatially evaluated by Lectin-IMC in relation to cell types and tissue microenvironments. MS-based glycoproteomics provides candidate carrier glycoproteins, which are subsequently incorporated into Lectin-IMC for higher-order evaluation of glycan–protein–cell type relationships. Lectin-IMC panel design is supported by a separate LMA-based probe-optimization module that assesses lectin–lectin interactions to support multiplex panel design. Spatially defined glyco-niches can further guide spatial (glyco)proteomic analysis, providing an iterative route back to candidate carrier discovery. (B) Representative histological and fluorescence images of left ventricular tissues from dilated cardiomyopathy (DCM) model and wild-type (WT) mice. Picrosirius red (PSR) staining shows collagen deposition and fibrotic regions. Fluorescence images show WFA-reactive glycans, WGA staining, and Hoechst nuclear staining. WFA signals are enriched in fibrotic regions of DCM hearts but do not completely overlap with PSR-positive collagen fibers, indicating selective localization of WFA-reactive glycans within fibrotic microenvironments. Boxed regions in whole-tissue images are shown at higher magnification. Scale bars, 500 μm.

## EXPERIMENTAL PROCEDURE

### Tissue samples

Mouse heart tissue sections used in this study were prepared from 24-week-old male 4C30 dilated cardiomyopathy (DCM) model mice and age-matched wild-type (WT) control mice, as described previously.^21^ The 4C30 mice were originally provided by Dr. Osamu Suzuki and Dr. Junichiro Matsuda from the Laboratory of Animal Models of the National Institutes of Biomedical Innovation, Health, and Nutrition (NIBIOHN, Osaka, Japan).^22^ Animal experiments for the preparation of these tissues were performed at the National Cerebral and Cardiovascular Center (NCVC) in accordance with relevant guidelines and regulations and were approved by the Institutional Animal Care and Use Committee of NCVC (approval no. 13040). Briefly, mice were anesthetized and perfused from the left ventricle with 0.1 M phosphate-buffered saline (PBS, pH 7.4), followed by 4% paraformaldehyde in PBS. Excised heart tissues were further fixed with 4% paraformaldehyde in PBS for 24 h at room temperature and embedded in paraffin. Horizontal FFPE cardiac sections were used for histological staining, lectin fluorescence staining, and Lectin-IMC analyses. The disease phenotype and cardiac remodeling parameters of the 4C30 and WT mice used for the preparation of these tissue sections were reported previously.^21^

Commercially available human FFPE heart tissue sections from a cardiomyopathy case were purchased from OriGene Technologies, Inc. (Rockville, MD, USA; Tissue FFPE Sections, Heart; CS808917). According to the supplier’s pathology verification, the tissue was derived from the heart and diagnosed as cardiomyopathy, with approximately 90% lesional tissue and 10% normal tissue, and no tumor or necrosis. The specimen was obtained from a 51-year-old female donor, and the diagnosis from the medical center pathology report was cardiomyopathy. The sections were stored at 4 °C according to the supplier’s instructions and used for Lectin-IMC analysis to evaluate the technical transferability of the optimized panel to human FFPE cardiomyopathy tissue. The specimens were supplied as anonymized research materials and contained no identifiable personal information. No human participants were recruited, enrolled, or contacted by the authors. According to the institutional policy of AIST, institutional ethics review was not required for the use of these commercially available specimens.

### Lectin–lectin interaction analysis using lectin microarray (LMA)

Lectin–lectin interaction analysis was performed using a lectin microarray according to previously described LMA procedures,^23^ with modifications to assess potential inter-lectin binding that could interfere with multiplex lectin staining. Fourteen representative lectins with distinct glycan-binding specificities were selected as applied lectin samples, and their binding profiles were analyzed on a lectin array chip containing 45 immobilized lectins (LecChip; GlycoTechnica, Yokohama, Japan). The abbreviations and carbohydrate specificities of the 45 immobilized lectins on the LecChip are summarized in Table S1. Information on the 14 applied lectin samples used for lectin–lectin interaction analysis is provided in Table S2. The complete lectin–lectin interaction data are provided in Table S3.

Briefly, each lectin sample was diluted in PBS containing 1% Triton X-100 (PBSTx), and 200 ng of lectin in 20 µL PBSTx was fluorescently labeled with 10 µg of Cy3-mono-reactive dye (Cytiva, Marlborough, MA, USA) for 1 h at room temperature in the dark. After labeling, 80 µL of probing buffer consisting of 25 mM Tris-HCl (pH 7.5), 137 mM NaCl, 2.7 mM KCl, 500 mM glycine, 1 mM CaCl₂, 1 mM MnCl₂, and 1% Triton X-100 was added, and the mixture was incubated for more than 30 min at room temperature in the dark to quench excess labeling reagent. The labeled lectin samples were applied to the LecChip at 60 µL per well, corresponding to 120 ng of lectin per well.

After application of the labeled lectins, the lectin array chips were incubated overnight at 20 °C in the dark. The chips were then washed three times with probing buffer and scanned using an evanescent-field fluorescence scanner, GlycoStation Reader 2300 (GSR2300; Emukk, Yokohama, Japan). All data were analyzed using GlycoStation Tools Pro Suite 1.5 software (GlycoTechnica). The net intensity was calculated by subtracting the background from the signal intensity. Data acquired with an exposure time of 532 ms in 1 × 1 binning mode were used to compare lectin– lectin interaction profiles after confirming that net intensities were below 40,000 for all spots.

### Histological and lectin fluorescence staining

Histological and lectin fluorescence staining procedures were performed based on previously described procedures^18^, with slight modifications. Tissue sections were deparaffinized three times with xylene for 5 min each and rehydrated through graded ethanol concentrations. Collagen deposition was evaluated by Picrosirius red (PSR) staining according to the manufacturer’s instructions.

For lectin fluorescence staining with WFA and WGA, antigen retrieval was performed by autoclaving the deparaffinized sections for 10 min at 110 °C in Target Retrieval Solution, pH 9.0 (Agilent Technologies, Santa Clara, CA, USA), diluted 10-fold with Milli-Q water. After antigen retrieval, the sections were washed with PBS and incubated with a streptavidin–biotin blocking kit (Vector Laboratories, Burlingame, CA, USA) to block endogenous biotin activity, followed by blocking with Carbo-Free Blocking Solution (Vector Laboratories) for 1 h at room temperature. The sections were then incubated with biotinylated WFA (2 µg/mL in PBS; Vector Laboratories) for 2 h at 20 °C, followed by incubation with Alexa Fluor 647-conjugated streptavidin (1 µg/mL in PBS; Thermo Fisher Scientific, Waltham, MA, USA) for 1 h at 20 °C in the dark. The sections were subsequently incubated with FITC-conjugated WGA (2 µg/mL in PBS; Vector Laboratories) and Hoechst 33342 (0.5 µg/mL in PBS; Dojindo, Kumamoto, Japan) for 1 h at 20 °C in the dark. After each incubation step, the sections were washed three times with PBS. The stained sections were mounted with ProLong Diamond Antifade Mountant (Thermo Fisher Scientific) and stored at 4 °C until imaging.

For validation of WFA reactivity, additional sections were subjected either to PNGase F digestion before the blocking steps or to competitive inhibition during WFA staining. For PNGase F treatment, after antigen retrieval and PBS washing, sections were incubated with PNGase F PRIME (25 mU/µL; N-Zyme Scientific, Doylestown, PA, USA) in reaction buffer at 37 °C for 16 h, followed by the blocking and lectin fluorescence staining procedures described above. For competitive inhibition, biotinylated WFA was preincubated with 200 mM GalNAc (Fujifilm Wako, Osaka, Japan) or 200 mM Gal (Fujifilm Wako) for 10 min at room temperature, and the mixture was then applied to the sections in place of untreated WFA. Subsequent detection with Alexa Fluor 647-conjugated streptavidin, WGA, and Hoechst 33342 was performed as described above.

PSR- and lectin fluorescence-stained sections were imaged in bright-field and fluorescence modes, respectively, using a NanoZoomer S60 slide scanner (Hamamatsu Photonics, Hamamatsu, Japan). Images were visualized and exported using NDP.view2 software (Hamamatsu Photonics).

### Preparation and chemical characterization of metal-labeled probes

Metal-labeled lectin probes were prepared by thiol modification of lectins followed by polymer-based metal conjugation, based on protein thiolation and mass cytometry antibody-labeling strategies described previously.^24,25^ The metal-conjugated probes and multiplex probe combinations used for Lectin-IMC are summarized in Table S4. Briefly, WFA, WGA, MAL-I, MAH, and SNA were adjusted to 0.5 mg/mL in PBS containing 5 mM EDTA, pH 8.0, and reacted with 0.8 mM Traut’s reagent (2-iminothiolane hydrochloride; Sigma-Aldrich, St. Louis, MO, USA) at room temperature for 1 h to introduce sulfhydryl groups into the lectins. The thiolated lectins were buffer-exchanged and washed with PBS using Amicon Ultra centrifugal filter units (10 kDa molecular weight cutoff; Millipore, Burlington, MA, USA). Lectin concentrations were determined by absorbance at 280 nm using an EzDrop micro-volume spectrophotometer (Blue-Ray Biotech, Taipei, Taiwan) with lectin-specific correction based on the A280 values of reference solutions of known protein concentration (Table S5).

The degree of thiol modification was determined using monobromobimane (mBBr)-based fluorescence labeling, as described previously.^26^ L-Cysteine (Fujifilm Wako) was used to generate a standard curve. Lectin samples or standards were mixed with mBBr and incubated for 15 min at room temperature in the dark. Fluorescence intensity was measured using an EnSpire multimode plate reader (PerkinElmer, Waltham, MA, USA) at excitation and emission wavelengths of 390 and 480 nm, respectively. The number of introduced thiol groups per lectin molecule was calculated from the measured thiol concentration and lectin concentration.

Thiolated lectins were conjugated to lanthanide-loaded maleimide-functionalized polymers through thiol–maleimide coupling to form stable thioether linkages, using the Maxpar X8 Multimetal Labeling Kit (Standard BioTools, South San Francisco, CA, USA) according to the manufacturer’s protocol, with modifications for lectin labeling. Metal-conjugated lectins were purified by centrifugal filtration and stored at 4 °C until use. Metal-labeled antibodies were prepared by conjugation with lanthanide-loaded polymers using the Maxpar X8 Multimetal Labeling Kit according to the manufacturer’s protocol.

The amount of metal incorporated into Dy162-labeled WFA was evaluated using the solution mode of the Hyperion Imaging System (Standard BioTools). A calibration curve was generated using dysprosium standard solution (Dy 1000; Fujifilm Wako) diluted in 2% nitric acid. The measured Dy concentration was converted to the Dy162 concentration based on the natural abundance of Dy162 (25.5%). The number of Dy162 atoms per WFA molecule was calculated from the Dy162 concentration and the WFA concentration determined by A280 measurement using the lectin-specific reference value (Table S5).

### Lectin-assisted imaging mass cytometry (Lectin-IMC)

The Lectin-IMC panel was designed to integrate complementary glycan, candidate carrier protein, and cell-type information required to characterize fibrosis-associated glyco-niches. WFA was included as the primary disease-associated glycan probe based on its previously demonstrated association with cardiac fibrogenic activity in the 4C30 DCM model.^21^ MAL-I, SNA, and MAH were included to provide complementary information on distinct sialylated glycan features, whereas WGA was used as a broad tissue-structural glycan probe. Selection of the multiplex lectin combination was further guided by the lectin–lectin interaction analysis described above. Cell-type and structural markers were selected to represent major components of the cardiac fibrotic microenvironment, including cardiomyocytes (MYL2), endothelial cells (CD31), mesenchymal/fibroblast-lineage cells (VIM), ACTA2-associated stromal and smooth muscle cells, and fibrotic extracellular matrix (COL1). Candidate ECM glycoproteins were selected from proteins previously identified in WFA-binding fractions by MS-based glycoproteomic analysis,^21^ considering their association with fibrotic ECM and the availability of antibodies compatible with FFPE multiplex imaging.

Lectin-IMC staining was performed according to the manufacturer’s recommended protocol for IMC sample preparation, with slight modifications for lectin detection. Briefly, FFPE tissue sections were deparaffinized and subjected to antigen retrieval by autoclaving for 10 min at 110 °C in Target Retrieval Solution, pH 9.0 (Agilent Technologies). After washing with PBS, the sections were blocked with Carbo-Free Blocking Solution (Vector Laboratories) for 1 h at room temperature. The sections were subsequently incubated with metal-conjugated lectins and antibodies diluted in Maxpar PBS (Standard BioTools). After washing, the sections were stained with DNA intercalator-Ir (Standard BioTools) for nuclear detection, washed with Maxpar Water (Standard BioTools), and air-dried at room temperature before IMC acquisition.

IMC data were acquired from selected regions of interest using a Hyperion Imaging System coupled with a Helios mass cytometer (Standard BioTools) at 200 Hz and 1 µm spatial resolution, as described previously^27^. Regions of interest were selected based on corresponding histological and lectin fluorescence images, with emphasis on PSR-positive fibrotic regions and WFA-reactive regions in this DCM mouse model.^21^

### Lectin-IMC data analysis

Lectin-IMC data were visualized using MCD Viewer software (Standard BioTools). Representative pseudo-colored images and channel overlays were generated in MCD Viewer and exported as image files for figure preparation. For WFA validation experiments, mean signal intensities of WFA and DNA in each region of interest (ROI) were obtained using MCD Viewer, and WFA intensity was normalized to the corresponding DNA signal before comparison with the untreated control.

Object-based colocalization maps were generated using the HALO image analysis software (Object Colocalization FL module v2.1.4; Indica Labs, Albuquerque, NM, USA). Four channels were used for colocalization analysis: ACTA2, VIM, POSTN, and WFA. Channel-specific object detection parameters, including intensity thresholds, contrast thresholds, and object-size ranges, were manually optimized for each marker based on signal intensity distributions and visual correspondence with marker-positive regions, and then applied consistently to comparable WT and DCM images without image-specific adjustment. Three object-based phenotypes were visualized using AND-positive criteria: ACTA2⁺VIM⁺ objects, ACTA2⁺VIM⁺WFA⁺ objects, and ACTA2⁺VIM⁺WFA⁺POSTN⁺ objects. The resulting colocalization maps were used to visualize the stepwise spatial refinement from ACTA2⁺VIM⁺ myofibroblast-like objects to WFA-associated and POSTN-associated higher-order spatial phenotypes in DCM and WT heart sections. Detailed channel-specific object detection parameters and phenotype definitions are provided in Table S6. Cell-based quantitative analysis was performed using QuPath software (version 0.6.0) ^28^. Cells were detected based on the nuclear DNA signal, and marker expression was quantified as mean intensity within the detected cell regions. Positivity thresholds for lectin and protein marker signals were determined based on signal intensity distributions, negative/background regions, and visual confirmation of marker-positive areas, and the same thresholds were applied consistently across comparable samples without image-specific adjustment. For cell-based analysis, cells were classified as WFA-positive when the mean WFA intensity within the detected cell region exceeded the predefined WFA threshold. Marker-positive cell populations were classified using the corresponding channel-specific thresholds.

For sub-ROI-based analysis, each 1 mm × 1 mm IMC image was automatically divided into 25 sub-ROIs. For distance-based analysis, WFA-positive regions were defined using a fixed WFA intensity threshold, and cells were classified according to their spatial relationship to WFA-positive regions. The distance from each marker-positive cell to the nearest WFA-positive region was calculated in QuPath.

Pixel-based spatial correspondence between WFA and candidate or control ECM proteins was quantified in a 1 mm × 1 mm IMC image acquired from a representative DCM tissue section. WFA-positive pixels were defined using the fixed intensity threshold established for WFA-positive region detection (≥8; Table S7). Protein-positive pixels were defined independently for each channel by global Otsu thresholding, and the resulting thresholds are provided in Table S8. Binary spatial overlap was expressed as the Dice coefficient, calculated as 2N_WFA ∩ protein_/(N_WFA_+N_protein_), where N denotes the number of positive pixels. Preferential protein localization to WFA-positive regions was quantified as the percentage of the total protein-channel signal intensity located within the WFA-positive mask. Mean signal enrichment was calculated as the ratio of the mean protein-channel intensity in WFA-positive pixels to that in WFA-negative pixels. The two intensity-based measures were calculated from the original protein-channel intensities without applying protein-positive pixel thresholds. All measurements were performed at the native spatial resolution of 1 μm per pixel.

Quality control was performed by visual inspection of cell detection results, tissue masks, and threshold-based marker-positive regions to exclude obvious segmentation artifacts and non-tissue/background areas. Detailed parameters for cell-based, sub-ROI-based, and distance-based QuPath analyses, including marker thresholds, cell-classification criteria, WFA-region definitions, and sub-ROI inclusion criteria, are provided in Table S7. Protein-positive pixel thresholds and the resulting pixel-based spatial correspondence measurements are summarized in Table S8. Quantitative values exported from QuPath were used for subsequent statistical analysis or descriptive comparison, as appropriate.

### Statistical analysis

A heat map for lectin–lectin interaction analysis was generated from log10-transformed net intensity values using GraphPad Prism 9 software (GraphPad Software, San Diego, CA, USA). The number of metal atoms per lectin molecule was calculated from three dilution series and presented as mean ± SD. Box-and-whisker plots and violin plots were generated using GraphPad Prism 9. Unless otherwise indicated, quantitative image-analysis data were analyzed at the ROI, sub-ROI, or cell level as specified in the figure legends. The pixel-based spatial correspondence analysis (Table S8) was performed on a single representative image and was used for descriptive comparison among candidate proteins; no inferential statistical testing was applied to these measurements. For sub-ROI- and cell-based spatial analyses, measurements were obtained from a single tissue specimen for each condition or specimen type. Sub-ROIs and individual cells therefore represent spatial subsamples or observations within the same biological specimen and were not treated as independent biological replicates. Mann–Whitney U-tests or Kruskal–Wallis tests followed by Dunn’s multiple comparisons tests were used as exploratory within-specimen comparisons of spatial distributions. The resulting P values are reported as nominal P values and should not be interpreted as evidence for population-level biological differences.

## RESULTS AND DISCUSSION

### Overview of the Multimodal Glycoprotein Analysis Framework

Figure 1A illustrates the multimodal glycoprotein analysis framework extended in this study to connect disease-associated glycan discovery, candidate carrier glycoprotein identification, and on-tissue spatial validation. The core workflow consists of stepwise discovery and spatial characterization of disease-associated glyco-niches. First, LMD-LMA tissue glycome mapping is used to identify disease-associated glycan alterations. Lectins corresponding to these glycan features are then incorporated into Lectin-IMC to evaluate their spatial relationships with major cellular and structural components of the tissue microenvironment. MS-based glycoproteomics provides candidate carrier glycoproteins associated with the disease-related glycan features, and antibodies against these candidates are subsequently incorporated into Lectin-IMC for higher-order evaluation of glycan–protein–cell type relationships. Within this framework, Lectin-IMC therefore functions as an on-tissue spatial validation and prioritization layer rather than as an isolated multiplex glycan-imaging method. LMA-based assessment of lectin–lectin interactions provides an additional probe-optimization module to support multiplex panel design. Together, these components enable disease-associated glycan features and MS-derived candidate glycoproteins to be reconnected within their spatial tissue context. In addition, glycan-, protein-, and cell-defined microenvironments identified by Lectin-IMC can serve as spatial entry points for subsequent image-guided proteomic or glycoproteomic analysis, providing a basis for an iterative discovery–validation workflow.

We first confirmed the distribution of fibrosis and WFA-reactive glycans in left ventricular tissues from dilated cardiomyopathy (DCM) model and wild-type (WT) mice using FFPE tissue sections corresponding to those analyzed in our previous study (Figure 1B). In this model, WFA-reactive asialo N-glycans were previously identified as fibrosis-associated glycan features.^21^ Picrosirius red (PSR) staining showed limited collagen deposition mainly in perivascular regions in WT hearts, whereas DCM hearts exhibited marked interstitial fibrosis. Immunofluorescence staining with Wisteria floribunda agglutinin (WFA), wheat germ agglutinin (WGA), and Hoechst revealed that WFA signals were enriched in fibrotic regions of DCM hearts. Notably, the WFA distribution did not completely overlap with collagen fibers visualized by PSR, indicating that WFA-reactive glycans represent a selective molecular feature within fibrotic microenvironments rather than a simple surrogate for total collagen deposition. These observations confirmed the reproducible enrichment of WFA-reactive glycans in fibrosis-associated regions and provided the histological basis for subsequent spatial validation within the multimodal framework.

### LMA-Guided Design and Characterization of Multiplex Lectin Probes for Lectin-IMC

To establish a multiplex lectin panel capable of providing complementary glycan information within WFA-defined fibrotic microenvironments, we first evaluated potential lectin–lectin interactions using LMA. Among the 45 lectins included in the LMA platform (Table S1), 14 representative lectins with distinct glycan-binding specificities were selected for comparison (Table S2). The heatmap of lectin–lectin binding signals revealed that several lectins, including AAL and ConA, showed broad inter-lectin binding profiles (Figure 2A; Table S3), indicating a greater potential for interference when combined with other lectins in multiplex staining. In contrast, WFA showed relatively limited interaction signals with MAL-I, SNA, MAH, and WGA, although a stronger interaction was observed with Jacalin. The final lectin panel was selected by considering both the biological information provided by each lectin and their compatibility for multiplex analysis. WFA served as the primary disease-associated glycan probe because WFA-reactive asialo N-glycans had previously been associated with cardiac fibrogenic activity in this DCM model.^21^ MAL-I, SNA, and MAH were incorporated to provide complementary information on distinct sialylated glycan features, whereas WGA was included as a broad tissue-structural glycan probe. Together with the LMA-based lectin–lectin interaction profiles, these complementary glycan specificities supported selection of WFA, MAL-I, SNA, MAH, and WGA as the five-lectin panel used for subsequent Lectin-IMC analysis (Table S4). Thus, this panel was designed not simply to enable multiplex staining, but to place the disease-associated WFA-reactive glycan feature within a broader spatial glycan context.

**Figure 2.**
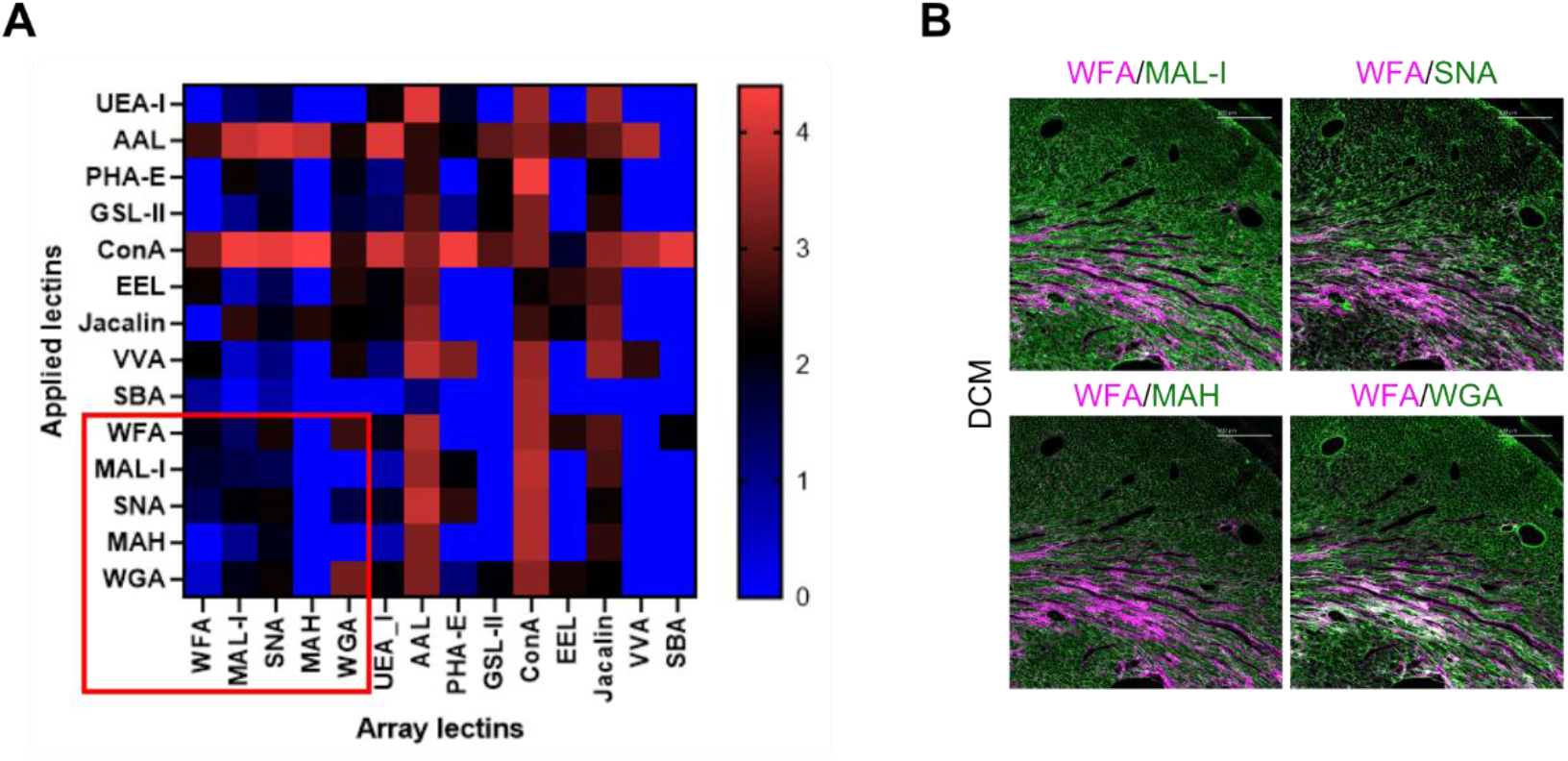
LMA-guided selection and characterization of lectin probes for multiplex Lectin-IMC. (A) Heatmap showing lectin–lectin interaction profiles among 14 representative lectins selected from the 45-lectin microarray platform. Signal intensities are shown as log10-transformed net intensity values. The red box indicates the five lectins selected for multiplex Lectin-IMC: WFA, MAL-I, SNA, MAH, and WGA. These lectins were selected based on their complementary glycan specificities and relatively low mutual interaction signals. (B) Representative two-channel views extracted from multiplex Lectin-IMC images, showing the spatial relationship between WFA and each selected lectin in cardiac tissue sections. WFA is shown in magenta, and MAL-I, SNA, MAH, or WGA is shown in green. MAL-I, SNA, and MAH exhibited spatial patterns distinct from WFA, whereas WGA showed partial overlap with WFA. Scale bars, 200 μm.

We next characterized the chemical modification of the metal-labeled lectin probes prepared for Lectin-IMC. Thiol incorporation after Traut’s reagent treatment was quantified as the number of introduced thiol groups per lectin molecule using an mBBr-based assay. Metal incorporation was evaluated specifically for Dy162-labeled WFA using the solution mode of the Hyperion Imaging System (Table S5). The numbers of introduced thiol groups per lectin molecule for the five selected lectins ranged from 0.085 to 0.367, confirming measurable thiol introduction under the present conditions. For WFA, solution-mode analysis indicated incorporation of approximately 11 Dy162 atoms per lectin molecule. These measurements provided chemical characterization of the metal-labeled lectin probes used for subsequent Lectin-IMC analysis.

To assess whether the selected lectins retained spatial distributions consistent with their known glycan-binding specificities on tissue sections, we performed multiplex Lectin-IMC imaging and compared the spatial distributions of WFA with MAL-I, SNA, MAH, or WGA using selected channel combinations (Figure 2B; Table S4). MAL-I, SNA, and MAH recognize distinct sialylated glycan features, whereas WFA preferentially recognizes terminal GalNAc-containing structures (Table S1). Consistent with these specificities, MAL-I, SNA, and MAH exhibited spatial patterns clearly distinct from WFA, with limited overlap in WFA-enriched fibrotic regions. In contrast, WGA showed a broader tissue distribution with partial spatial overlap with WFA. These results indicate that the selected lectins retained spatial distributions broadly consistent with their respective glycan-binding specificities after metal labeling and multiplex tissue staining, supporting their combined use as a multiplex panel for Lectin-IMC.

### On-Tissue Validation of Glycan-Dependent Lectin Binding by Lectin-IMC

We next evaluated whether metal labeling and IMC detection preserved glycan-dependent lectin binding on tissue sections. Because WFA-reactive asialo N-glycans had previously been identified as fibrosis-associated glycan features in this DCM model, WFA was used as the representative disease-associated lectin probe for analytical validation. WFA binding was evaluated by competitive inhibition and PNGase F-mediated N-glycan removal using adjacent cardiac tissue sections. Under control conditions, WFA signals were detected in fibrotic regions by both conventional immunofluorescence (IF) and Lectin-IMC (Figure 3). In the presence of competitive sugars, WFA signals were attenuated, with GalNAc producing a stronger inhibitory effect than Gal. PNGase F treatment also markedly reduced the WFA signals, indicating that a substantial fraction of the detected signal depended on N-glycans. Importantly, these signal changes were observed consistently by both IF and Lectin-IMC, demonstrating that the metal-labeling and IMC-detection workflow preserved glycan-dependent WFA binding on tissue sections.

**Figure 3.**
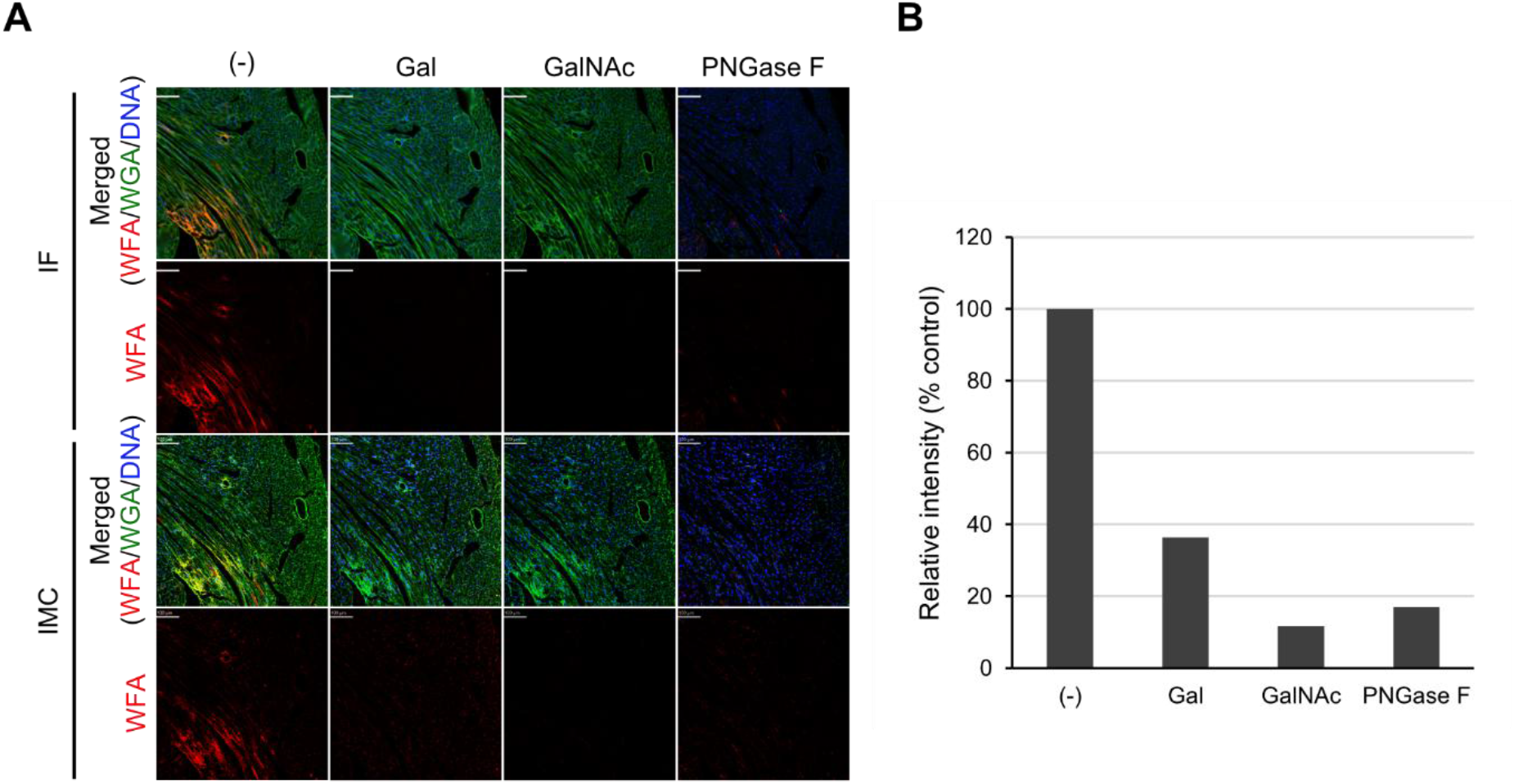
On-tissue validation of glycan-dependent WFA binding by Lectin-IMC. (A) Representative images comparing WFA staining by conventional immunofluorescence (IF) and Lectin-IMC in cardiac tissue sections under control conditions, competitive inhibition with galactose (Gal) or N-acetylgalactosamine (GalNAc), and PNGase F treatment. Merged images show WFA, WGA, and DNA/Hoechst staining; corresponding WFA single-channel images are shown below each merged image. WFA signals were reduced by competitive sugars and PNGase F treatment in both IF and Lectin-IMC. (B) Quantification of WFA signal intensity from Lectin-IMC images. Mean WFA intensity was normalized to the DNA signal and expressed as relative intensity compared with the untreated control condition. Gal, GalNAc, and PNGase F treatment reduced WFA signals, supporting glycan-dependent detection of WFA-reactive glycans by Lectin-IMC. Scale bars, 100 μm.

We further quantified WFA signal intensity from the Lectin-IMC data (Figure 3B). Mean WFA intensity was calculated for each region of interest and normalized to the DNA signal, and relative intensity was expressed as a percentage of the untreated control. Competitive inhibition with Gal and GalNAc reduced the WFA signal to approximately 36% and 12% of the control level, respectively, whereas PNGase F treatment reduced the signal to approximately 17%. These quantitative results further support the analytical specificity of metal-labeled WFA detection and establish the basis for subsequent spatial analysis of fibrosis-associated WFA-reactive glycans by Lectin-IMC.

### Cell-Type–Resolved Mapping of WFA-Reactive Glycans in Fibrotic Microenvironments

Having established glycan-dependent detection of WFA-reactive signals, we next examined their spatial relationships with major cellular and structural components of the cardiac fibrotic microenvironment. The marker panel was designed as a focused set representing cardiomyocytes (MYL2), endothelial cells (CD31), mesenchymal/fibroblast-lineage cells (VIM), ACTA2-associated stromal and smooth muscle cells, and fibrotic extracellular matrix (COL1). A shows representative Lectin-IMC images combining these markers with the five-lectin panel (Table S4). The lectin channels exhibited distinct spatial distributions, with WFA showing characteristic enrichment in fibrotic regions. The marker panel showed accumulation of VIM- and ACTA2- positive cells together with COL1-positive ECM within these regions. Thus, Lectin-IMC enabled disease-associated glycan signals to be evaluated in relation to multiple cellular and structural features within the same tissue context.

We next evaluated the relationship between WFA-reactive glycans and individual cellular markers. WFA signals did not exhibit clear global overlap with ACTA2, MYL2, CD31, or VIM alone (Figure S1). Because our previous glycoproteomic analysis suggested that WFA-reactive glycans were predominantly associated with ECM glycoproteins, we focused on fibroblast-lineage populations within fibrotic regions, particularly VIM-positive cells and the ACTA2⁺VIM⁺ myofibroblast-like population. WFA-enriched regions spatially corresponded to areas containing VIM-positive cell accumulations, with partial association with ACTA2-positive regions (Figure 4B). Consistently, ACTA2⁺VIM⁺ cells associated with WFA signals were preferentially observed along fibrotic areas (Figure 4C).

**Figure 4.**
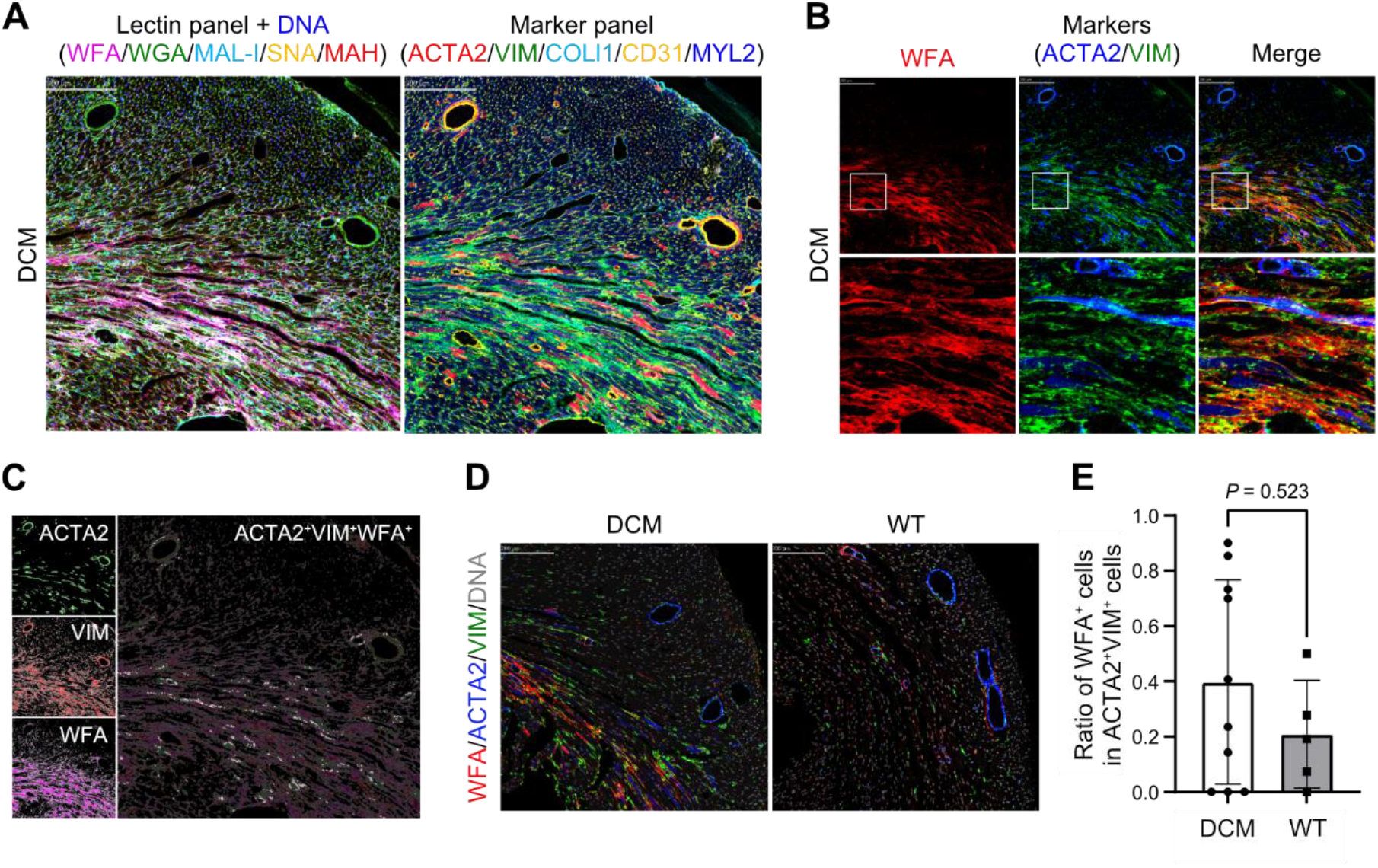
Cell-type–resolved mapping of WFA-reactive glycans in fibrotic microenvironments. (A) Representative Lectin-IMC images of corresponding fibrotic regions stained with a lectin panel (WFA, WGA, MAL-I, SNA, MAH, and DNA) and a marker panel (VIM, COL1, CD31, ACTA2, and MYL2). WFA was enriched in fibrotic regions, whereas the marker panel revealed accumulation of VIM- and ACTA2-positive cells together with COL1-positive ECM. (B) Spatial relationship between WFA and fibroblast/myofibroblast markers. Single-channel and merged images of WFA, ACTA2, and VIM are shown, with boxed regions enlarged below. WFA-enriched regions spatially corresponded to areas containing VIM-positive cells and partially overlapped with ACTA2-positive regions. (C) Visualization of ACTA2⁺VIM⁺ cells associated with WFA signals. Left panels show individual distributions of ACTA2, VIM, and WFA, and the right panel shows the spatial distribution of ACTA2⁺VIM⁺ cells classified as WFA-associated based on the image-analysis criteria. These WFA-associated ACTA2⁺VIM⁺ cells were preferentially observed along fibrotic regions. (D) Representative Lectin-IMC images from DCM and wild-type (WT) left ventricular tissues stained for WFA, VIM, ACTA2, and DNA. (E) Quantification of the proportion of ACTA2⁺VIM⁺ cells classified as WFA-positive in DCM and WT tissues. Although the DCM tissue showed a trend toward an increased proportion of WFA-positive cells, the difference was not statistically significant (Mann–Whitney U-test). Data points represent sub-ROIs; bars indicate mean ± SEM. Scale bars, 200 μm.

To quantitatively examine this relationship, corresponding left ventricular regions from DCM and WT hearts were analyzed under identical conditions (Figure 4D), and the proportion of ACTA2⁺VIM⁺ cells classified as WFA-positive was calculated using the QuPath-based cell-classification and sub-ROI analysis workflow (Figure S2 and Table S7). The analyzed DCM tissue showed a trend toward a higher proportion of ACTA2⁺VIM⁺ cells classified as WFA-positive than the WT tissue; however, this difference was not statistically significant under the present analysis conditions (Figure 4E).

Together, these results indicate that WFA-reactive glycans are spatially enriched within fibrotic microenvironments containing ACTA2⁺VIM⁺ myofibroblast-like cells but cannot be unambiguously assigned to a specific cell population based on glycan–cell type mapping alone. The incomplete spatial correspondence and lack of a significant difference in the present sub-ROI-based comparison therefore expose an analytical limitation of glycan localization alone and provide the rationale for incorporating candidate carrier glycoprotein information in the next stage of the framework.

### Spatial Integration of WFA-Reactive Glycans and Candidate Carrier Glycoproteins Characterizes a Fibrosis-Associated Glyco-Niche

The cell-type–resolved analysis indicated that WFA-reactive glycans are preferentially distributed within fibrotic regions containing ACTA2⁺VIM⁺ myofibroblast-like cells. However, spatial glycan and cell-type information alone cannot determine which molecular components contribute to these signals. We therefore incorporated MS-derived candidate carrier glycoproteins into Lectin-IMC to evaluate glycan–protein–cell type relationships within the same tissue context. Based on our previous glycoproteomic analysis,^21^ ECM glycoproteins detected at the glycopeptide level in WFA-binding fractions from DCM hearts were considered candidate carriers of WFA-reactive N-glycans. Six candidate proteins for which antibodies applicable to tissue staining were available were selected for spatial evaluation: BGN, COL6A6, POSTN, HSPG2, LAMA4, and LAMC1. The distributions of the six candidate proteins were compared with that of WFA in the same DCM tissue section (Figure 5A, Figure S3, and Table S4). The evaluated proteins exhibited distinct localization patterns within fibrotic regions. To objectively assess their spatial correspondence, we calculated the Dice coefficient, the proportion of total protein-channel signal located within WFA-positive regions, and the enrichment of mean protein-channel signal in WFA-positive relative to WFA-negative regions (Table S8). Among the candidate proteins, POSTN showed the highest Dice coefficient (0.410), the largest proportion of protein signal within WFA-positive regions (53.1%), and the greatest mean signal enrichment in WFA-positive regions (8.04-fold). COL6A6 showed the second-highest Dice coefficient among the candidates (0.355), whereas the other candidate proteins showed lower spatial overlap and signal enrichment. These quantitative results were consistent with the merged images and supported prioritization of POSTN as the candidate showing the strongest spatial correspondence with WFA-positive tissue structures in the analyzed IMC image.

**Figure 5.**
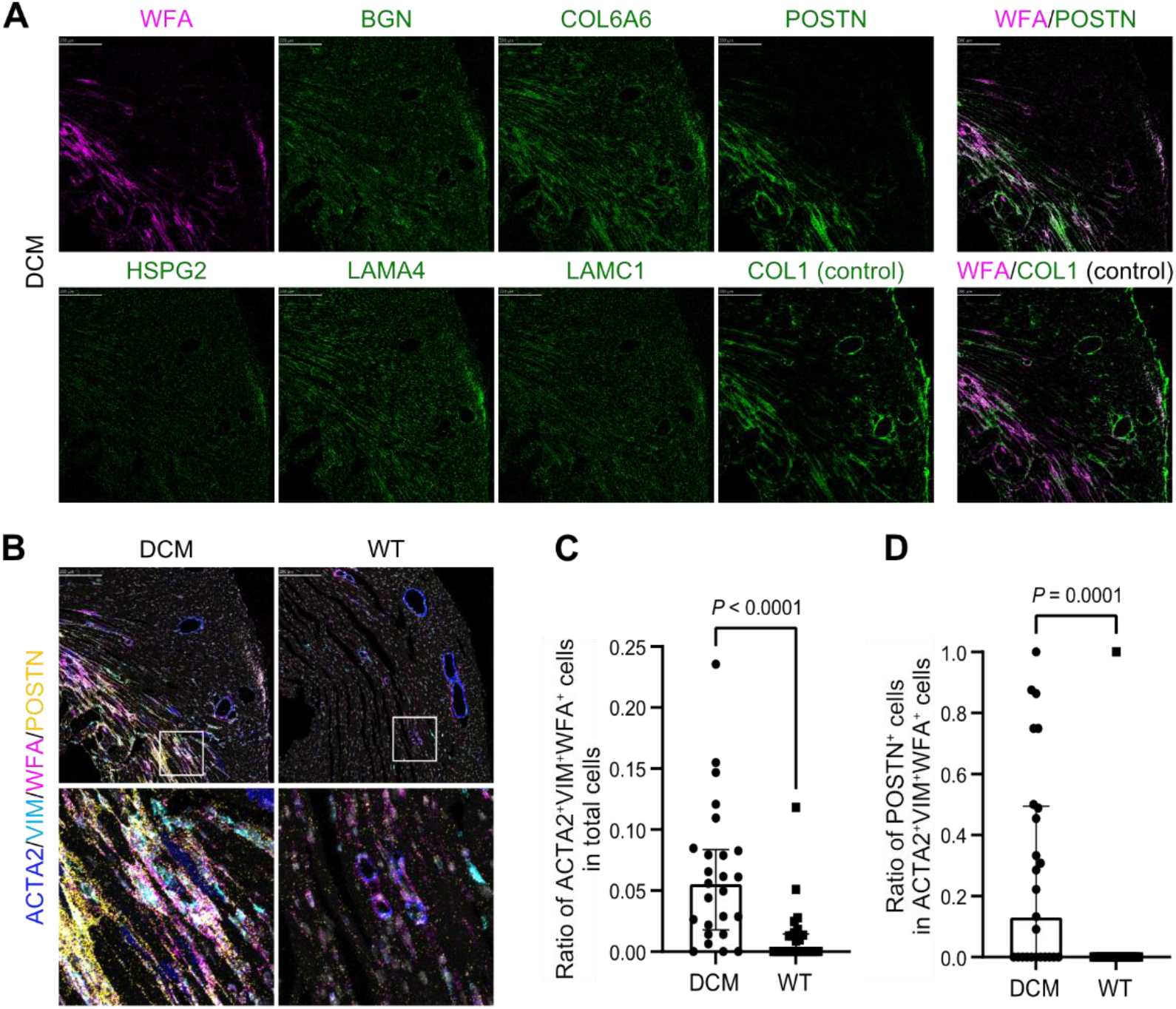
Spatial integration of WFA-reactive glycans and candidate carrier glycoproteins characterizes a fibrosis-associated glyco-niche. (A) Representative Lectin-IMC images comparing the spatial distributions of WFA and candidate ECM glycoproteins detected at the glycopeptide level in WFA-binding fractions by previous glycoproteomic analysis. Candidate proteins included BGN, COL6A6, POSTN, HSPG2, LAMA4, and LAMC1. COL1 was included as a structural ECM control. POSTN showed the strongest spatial correspondence with WFA-enriched regions among the tested candidate proteins, whereas other ECM proteins exhibited broader or partially distinct distributions, as supported by the quantitative pixel-based analysis summarized in Table S8. Corresponding WFA/protein merged images are shown in Figure S3. (B) Representative four-channel Lectin-IMC images of WFA, POSTN, ACTA2, and VIM in DCM and WT heart tissues, with boxed regions shown at higher magnification below. POSTN signals were enriched in ACTA2⁺VIM⁺WFA⁺ fibrotic regions in DCM hearts, whereas such spatial association was limited in WT hearts. (C) Quantification of the proportion of ACTA2⁺VIM⁺WFA⁺ cells among total cells across spatial sub-ROIs from representative DCM and WT tissue sections. (D) Quantification of the proportion of POSTN-positive cells within the ACTA2⁺VIM⁺WFA⁺ cell population across spatial sub-ROIs. Each data point represents one spatial sub-ROI derived from a single tissue specimen per condition and therefore does not represent an independent biological replicate. Mann–Whitney U-tests were used as exploratory comparisons of the sub-ROI distributions, and nominal P values are shown. These comparisons are intended to describe within-specimen spatial patterns rather than population-level biological differences. Bars indicate mean ± SEM. Scale bars, 200 μm.

We therefore focused on POSTN for higher-order spatial analysis together with WFA, ACTA2, and VIM. Four-channel imaging showed that POSTN signals were enriched in ACTA2⁺VIM⁺ WFA⁺ regions in DCM hearts, whereas this spatial association was limited in WT hearts (Figure 5B). Magnified views further showed POSTN along WFA-enriched fibrotic structures containing ACTA2⁺VIM⁺ cells, spatially prioritizing POSTN as a candidate component of the WFA-defined glyco-niche. HALO-based object colocalization analysis further visualized this hierarchical spatial relationship: ACTA2⁺VIM⁺ objects were broadly distributed within fibrotic regions, whereas ACTA2⁺VIM⁺WFA⁺POSTN⁺ objects were preferentially localized within WFA-enriched structures in DCM hearts and were rarely observed in WT hearts (Figure S4 and Table S6).

Quantitative analysis of the corresponding spatial phenotypes further supported this pattern. The distribution of ACTA2⁺VIM⁺WFA⁺ cell proportions across the analyzed sub-ROIs differed between the representative DCM and WT tissue sections (Figure 5C; nominal P < 0.0001, Mann– Whitney U-test). The proportion of POSTN-positive cells within the ACTA2⁺VIM⁺WFA⁺ population was also higher across sub-ROIs in the representative DCM tissue section than in the WT tissue section (Figure 5D; nominal P = 0.0001, Mann–Whitney U-test). These within-specimen analyses support the spatial association of POSTN with WFA-associated, ACTA2⁺VIM⁺-rich fibrotic regions in the analyzed DCM tissue.

Taken together, these results demonstrate how integration of MS-derived candidate glycoprotein information with Lectin-IMC extends spatial analysis from glycan–cell type relationships to glycan–protein–cell type relationships. In the DCM fibrotic microenvironment, this approach spatially prioritized POSTN among the tested MS-derived candidate carrier glycoproteins as a component associated with the WFA-defined glyco-niche.

### Applicability of the Lectin-IMC Panel to a Human FFPE Cardiomyopathy Specimen

We next evaluated the technical transferability of the Lectin-IMC panel optimized in the mouse DCM model to a human FFPE cardiomyopathy specimen. Using the same lectin and antibody panel, we compared the spatial distributions of WFA-reactive glycans, complementary lectin signals, and major cardiac cell-type markers in mouse DCM and human cardiomyopathy tissue sections (Figure 6A and Table S4). In both specimen types, WFA signals were detected in restricted tissue regions corresponding to fibrotic or fibrosis-like areas. The multiplex panel also enabled simultaneous visualization of glycan signals and major cellular and structural markers in the human FFPE specimen, although signal intensities and detailed spatial distributions differed between the mouse and human tissues. These observations demonstrate the technical applicability of the multiplex staining and imaging workflow to human FFPE cardiomyopathy tissue.

**Figure 6.**
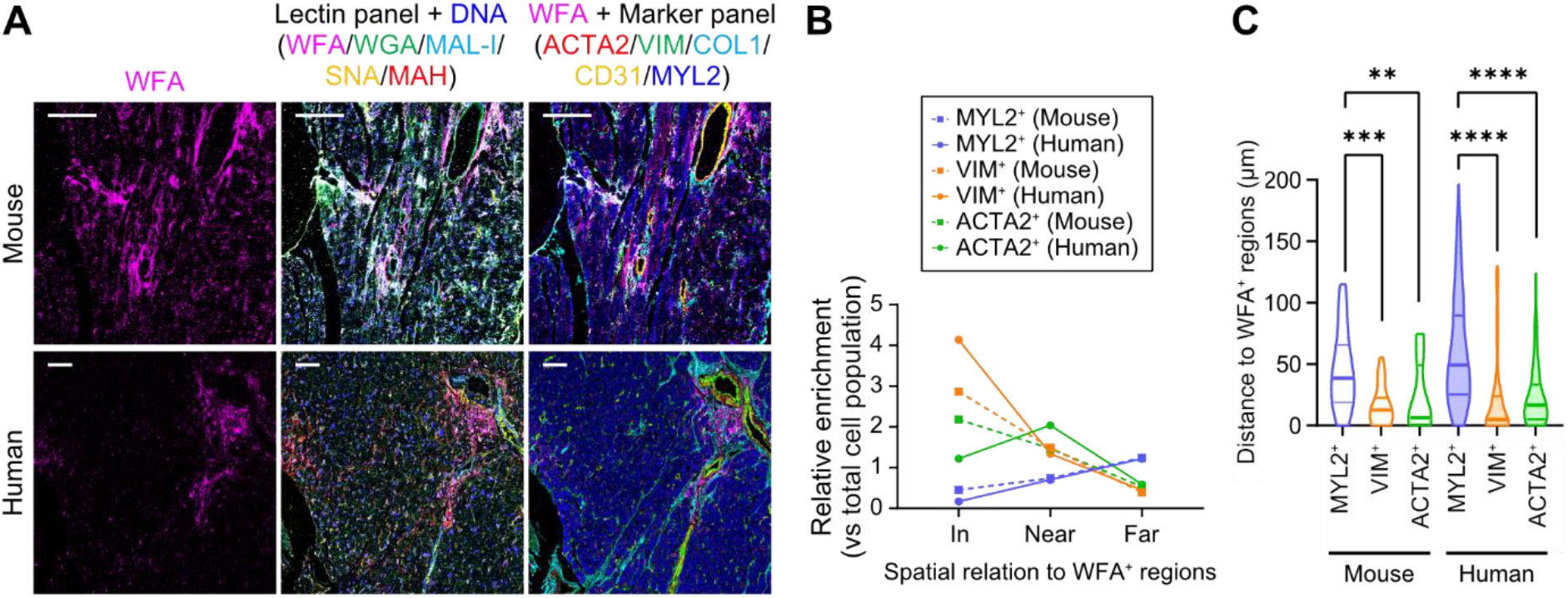
Applicability of the Lectin-IMC panel to a human FFPE cardiomyopathy specimen. (A) Representative Lectin-IMC images comparing mouse DCM and human cardiomyopathy tissue sections stained with the same lectin and antibody panel. WFA single-channel images, merged lectin panel images, and merged images combining WFA with cell-type markers are shown. The lectin panel included WFA, WGA, MAL-I, SNA, MAH, and DNA, and the marker panel included VIM, COL1, CD31, ACTA2, and MYL2. WFA signals were detected in restricted fibrotic or fibrosis-like regions in both mouse and human tissues, demonstrating applicability of the panel to human FFPE cardiomyopathy tissue. (B) Relative enrichment of MYL2⁺, VIM⁺, and ACTA2⁺ cells according to their spatial relationship to WFA-positive regions. Cells were classified as located within WFA-positive regions, near WFA-positive regions, or far from WFA-positive regions. VIM⁺ cells were enriched within WFA-positive regions in both mouse and human tissues, whereas MYL2⁺ cardiomyocytes were relatively depleted within WFA-positive regions and increased at greater distances. ACTA2⁺ cells showed an intermediate distribution. (C) Distance distributions from MYL2⁺, VIM⁺, and ACTA2⁺ cells to the nearest WFA-positive region within representative mouse DCM and human cardiomyopathy tissue sections. Statistical comparisons among cell-type-specific distance distributions within each specimen were performed using the Kruskal–Wallis test followed by Dunn’s multiple comparisons test. Individual cells represent spatial observations within a single tissue specimen and are not independent biological replicates. The resulting nominal P values are provided as exploratory measures of differences among within-specimen spatial distributions and are not intended for population-level inference. **P < 0.01, ***P < 0.001, ****P < 0.0001. Scale bars, 100 μm.

To characterize the spatial organization of cell types relative to WFA-positive regions, cells were classified according to their localization relative to WFA-positive areas, and the relative enrichment of MYL2⁺, VIM⁺, and ACTA2⁺ cells was calculated (Figure 6B and Table S7). Within the representative specimens examined, VIM⁺ cells were enriched within WFA-positive regions and decreased with increasing distance from these regions. In contrast, MYL2⁺ cardiomyocytes were relatively depleted within WFA-positive regions and tended to increase in regions farther from WFA-positive areas, whereas ACTA2⁺ cells showed an intermediate pattern.

We further analyzed the distance of each cell type from the nearest WFA-positive region (Figure 6C and Table S7). Within the representative mouse DCM and human cardiomyopathy sections, VIM⁺ cells were generally located closer to WFA-positive regions than MYL2⁺ cells, whereas ACTA2⁺ cells showed an intermediate distribution (Figure 6C). These distance-based analyses illustrate that the Lectin-IMC workflow can quantify cell-type organization around glycan-defined regions in both mouse and human FFPE tissues. Collectively, these results support the technical transferability of the mouse-optimized Lectin-IMC panel to a human cardiomyopathy specimen and illustrate its potential for comparative spatial analysis across specimen types.

### Implications for Spatial Glycopathology and Glycoprotein-Targeted Discovery

The present study demonstrates the value of interpreting disease-associated glycan signals together with candidate carrier glycoprotein and cell-type information. WFA-reactive glycans were preferentially distributed within fibrotic regions enriched in ACTA2⁺VIM⁺ myofibroblast-like cells, whereas glycan signals alone showed incomplete correspondence with individual cell markers and were insufficient to define a specific cellular or molecular substrate (Figure 4). This illustrates a general challenge in spatial glycan analysis: a lectin-detected glycan motif can be present on multiple glycoproteins distributed across different cellular and extracellular compartments. Incorporation of MS-derived candidate carrier glycoproteins into Lectin-IMC addressed this limitation by extending the analysis from glycan–cell type relationships to glycan– protein–cell type relationships. Among the tested candidates, POSTN showed the strongest spatial correspondence with WFA-positive regions in the pixel-based quantitative analysis and was further associated with WFA-reactive glycans in ACTA2⁺VIM⁺-rich fibrotic regions and was thereby spatially prioritized as a candidate component of the WFA-defined glyco-niche (Figure 5). Application of the same panel to a human FFPE cardiomyopathy specimen further demonstrated the technical transferability of this analytical workflow (Figure 6).

Our previous LMD-LMA–Lectin-LVSEM workflow provided complementary structural information by resolving lectin-reactive glycans within fibrotic ECM architecture at high spatial resolution.^18^ The present framework extends this strategy by reconnecting disease-associated glycan features with cellular context and MS-derived candidate glycoproteins in situ. In this context, Lectin-IMC serves not simply as a multiplex lectin-imaging platform but as a spatial validation and prioritization layer linking upstream glycan and glycoprotein discovery with tissue-level molecular organization.

LMA-based assessment of lectin–lectin interactions provides an additional analytical design element for constructing multiplex lectin panels. Importantly, the final panel was selected by considering both potential inter-lectin interference and the complementary biological information provided by individual probes. More broadly, the multimodal framework can operate iteratively rather than as a one-directional sequence. Glycan-, protein-, and cell-informed microenvironments identified by Lectin-IMC can define regions or cell populations for subsequent image-guided proteomic or glycoproteomic interrogation. Thus, upstream molecular discovery can inform spatial validation, while spatially defined niches can in turn guide deeper molecular analysis.

### Limitations and Future Directions

Several limitations should be considered when interpreting this study. First, although Lectin-IMC enables glycan and protein signals to be visualized within the same tissue context, spatial correspondence does not demonstrate that a particular protein directly carries the lectin-detected glycan. Our previous glycoproteomic analysis identified multiple ECM proteins, including POSTN, in association with WFA-reactive glycans. The spatial correspondence observed here therefore supports prioritization of POSTN as a candidate component of the WFA-defined glyco-niche, but does not establish direct carriage of the WFA-reactive glycan by POSTN in situ. Direct glycoform-level confirmation will require orthogonal molecular approaches.

Second, the quantitative spatial analyses were performed on representative tissue specimens, and the pixel-based comparison of candidate ECM proteins was based on a single representative DCM IMC image. These analyses were therefore intended to characterize within-specimen spatial relationships and prioritize candidate proteins rather than to establish population-level biological differences. Validation in independent biological specimens will be required to assess the reproducibility of these spatial associations.

An important future direction is to connect Lectin-IMC-defined glyco-niches with image-guided spatial proteomics and glycoproteomics. Deep visual proteomics has demonstrated that microscopy-based phenotyping, image-guided selection, laser microdissection, and ultrasensitive MS can be integrated to profile proteomes of spatially defined cell populations while preserving tissue context.^10^ In this setting, Lectin-IMC could provide glycan-informed spatial phenotypes that complement morphology- or protein-based image guidance. Glycan- and protein-defined microenvironments identified by Lectin-IMC could therefore serve as spatial entry points for downstream region- or cell population-selective proteomics and potentially glycoproteomics, creating an iterative workflow in which spatial validation guides the next round of molecular discovery. Further evaluation in additional disease models, organs, and clinical specimens will be required to determine the broader generalizability of the framework.

## CONCLUSIONS

We extended a multimodal glycoprotein analysis framework by incorporating Lectin-IMC as an on-tissue spatial validation and prioritization layer linking disease-associated glycan discovery, candidate carrier glycoprotein identification, and glycan–protein–cell type analysis. Multiplex panel design was supported by LMA-based assessment of lectin–lectin interactions and biologically informed selection of lectin, cell-type, structural, and candidate carrier protein markers. Using fibrosis-associated WFA-reactive glycans in a DCM model, we validated glycan-dependent detection by metal-labeled WFA and spatially related these glycans to ACTA2⁺VIM⁺ myofibroblast-like populations. Pixel-based comparison of six MS-derived candidate ECM glycoproteins showed that POSTN had the strongest spatial correspondence with WFA-positive regions, and higher-order Lectin-IMC analysis further associated POSTN with ACTA2⁺VIM⁺WFA⁺ fibrotic microenvironments. These findings spatially prioritize POSTN as a candidate component of the WFA-defined glyco-niche without establishing direct glycoform-level association in situ. The optimized Lectin-IMC panel was also technically transferable to a human FFPE cardiomyopathy specimen. By preserving tissue architecture while simultaneously visualizing glycan, protein, and cell-type information, Lectin-IMC provides intuitive spatial interpretability of disease-associated glyco-niches. More broadly, spatially defined glyco-niches identified by Lectin-IMC may provide entry points for subsequent proteomic or glycoproteomic analysis, allowing the framework to operate as an iterative cycle linking molecular discovery, spatial validation, and candidate prioritization. This framework provides a practical strategy for integrating glycan information into spatial analysis of disease-associated glycoproteins within tissue microenvironments.

## ASSOCIATED CONTENT

### Supporting Information

The following files are available free of charge.

Additional Lectin-IMC images, QuPath-based image-analysis workflow, candidate extracellular matrix glycoprotein comparisons, and HALO object colocalization maps (PDF). Lectin and probe information, lectin–lectin interaction data, multiplex Lectin-IMC panel compositions, lectin-probe characterization, and HALO and QuPath image-analysis parameters (XLSX).

## AUTHOR INFORMATION

### Corresponding Author

Chiaki Nagai-Okatani. Cellular and Molecular Biotechnology Research Institute, National Institute of Advanced Industrial Science and Technology (AIST), 1-1-1 Higashi, Tsukuba, Ibaraki 305-8565, Japan.

### Author Contributions

Conceptualization: C.N.-O. and A.K.; Methodology: C.N.-O. and A.K.; Formal analysis: C.N.-O., U.H., P.B., and K.M.; Funding acquisition: C.N.-O.; Writing – original draft: C.N.-O.; Writing – review and editing: C.N.-O., U.H., P.B., K.M., and A.K. The manuscript was written through contributions of all authors. All authors have given approval to the final version of the manuscript.

### Notes

The authors declare no competing financial interest.

## ACKNOWLEDGMENTS

We thank Dr. Hiroyuki Kaji (Nagoya University) for valuable advice on the Lectin-IMC concept. We also thank Dr. Mitsuhiro Nishigori and Dr. Naoto Minamino for their contributions at the National Cerebral and Cardiovascular Center (NCVC) to animal care, tissue collection, and preparation of the FFPE heart tissue specimens used in this study. Figure 1A and the Table of Contents graphic were created with BioRender.com. Generative artificial intelligence was used during manuscript preparation to assist with language editing and improving clarity and organization. All scientific content, interpretations, and final text were reviewed and approved by the authors.

