## Supporting Information for "Lectin-Assisted Imaging Mass Cytometry (Lectin-IMC) Enables Spatial Validation of Disease-Associated Glyco-Niches within a Multimodal Glycoprotein Analysis Framework"

### Table of Contents

#### 1. Supporting Figures

Figure S1. Spatial relationship between WFA-reactive glycans and individual cardiac cell markers.

Figure S2. QuPath-based cell detection, marker thresholding, and sub-ROI division for Lectin-IMC quantification.

Figure S3. Spatial comparison of WFA-reactive glycans and candidate extracellular matrix glycoproteins.

Figure S4. HALO object colocalization maps of ACTA2, VIM, WFA, and POSTN in DCM and WT hearts.

#### 2. Supporting Tables (separate XLSX file)

Table S1. Abbreviations and carbohydrate specificities of 45 lectins immobilized on the LMA.

Table S2. Applied lectin samples used for lectin–lectin interaction analysis.

Table S3. Net intensity data of LMA-based lectin–lectin interaction analysis.

Table S4. Metal-conjugated probes and probe combinations used in Lectin-IMC experiments.

Table S5. Chemical characterization of thiolated and metal-labeled lectin probes.

Table S6. Parameters used for HALO-based object colocalization analysis.

Table S7. QuPath image analysis parameters.

Table S8. Pixel-based spatial correspondence between WFA and selected ECM proteins.

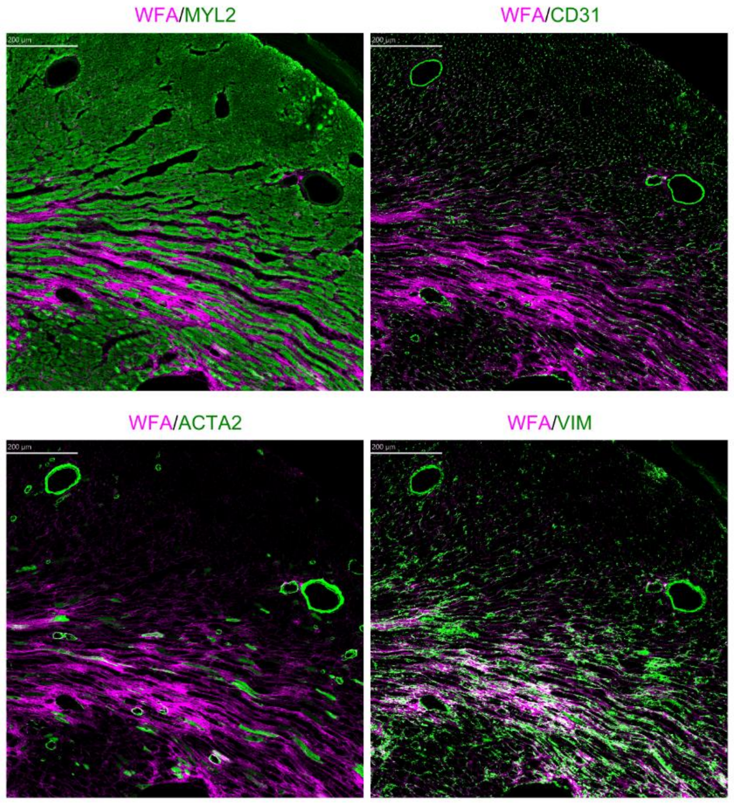


**Figure S1. Spatial relationship between WFA-reactive glycans and individual cardiac cell markers.** Representative Lectin-IMC images showing the spatial relationships between WFA and MYL2, CD31, ACTA2, or VIM in a DCM heart section. WFA is shown in magenta, and each protein marker is shown in green. MYL2, CD31, ACTA2, and VIM were used to visualize cardiomyocytes, endothelial cells, smooth muscle/myofibroblast-related cells, and mesenchymal/fibroblast-lineage cells, respectively. WFA showed limited spatial correspondence with MYL2, CD31, and ACTA2, whereas partial correspondence with VIM was observed in WFA-enriched fibrotic regions. Display ranges for each channel are indicated in the figure. Scale bars, 200 μm.

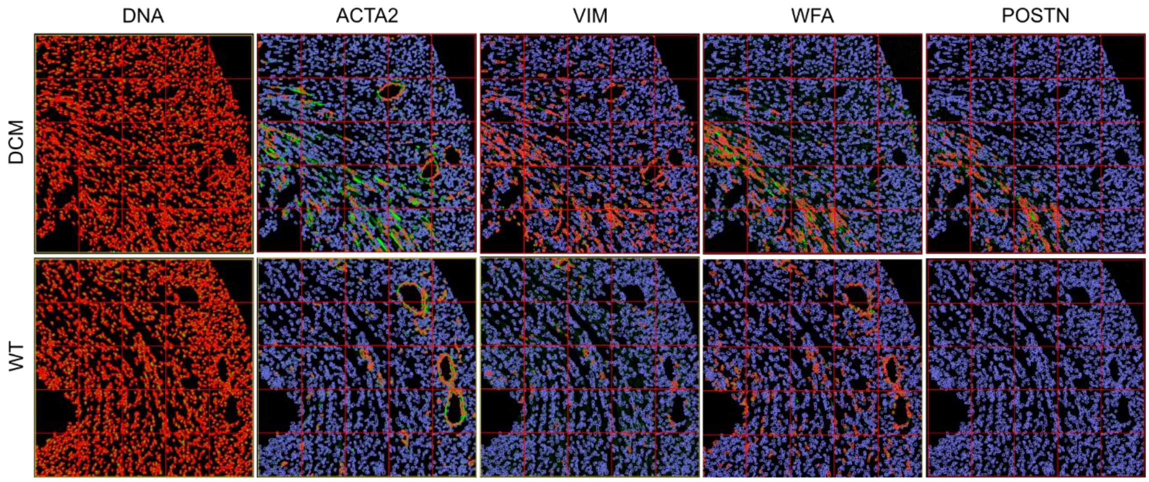


**Figure S2. QuPath-based cell detection, marker thresholding, and sub-ROI division for Lectin-IMC quantification.** Representative QuPath analysis outputs from DCM and WT heart sections are shown for DNA, ACTA2, VIM, WFA, and POSTN. In the DNA panels, nuclei are shown in green and detected cells are indicated in red. In the ACTA2, VIM, WFA, and POSTN panels, nuclei are shown in blue, detected cells are indicated in red, and the corresponding threshold-based marker-positive signals are shown in green. Each 1 mm × 1 mm IMC image was divided into 25 sub-ROIs of 200 μm × 200 μm. For analysis of WFA positivity within the $VIM^{+}ACTA2^{+}$ cell population, only sub-ROIs containing at least 10 $VIM^{+}ACTA2^{+}$ cells were included. Detailed QuPath analysis parameters, including marker thresholds and sub-ROI inclusion criteria, are provided in Table S7.

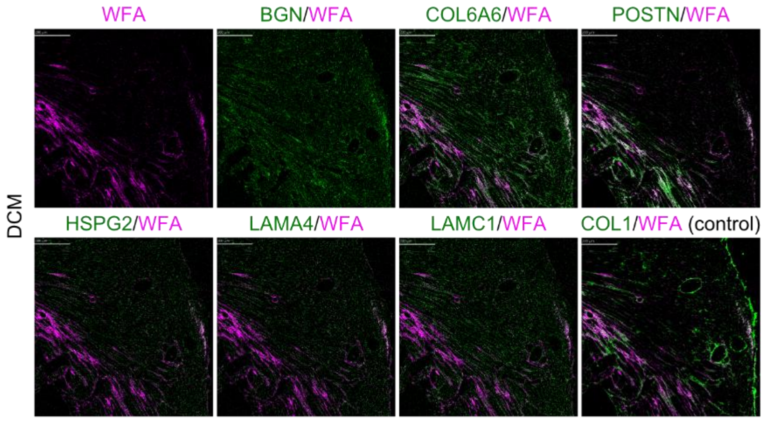


**Figure S3. Spatial comparison of WFA-reactive glycans and candidate extracellular matrix glycoproteins.** Representative Lectin-IMC images showing WFA alone and its spatial relationships with candidate ECM glycoproteins detected at the glycopeptide level in WFA-binding fractions by MS-based glycoproteomic analysis. WFA is shown in magenta, and each candidate glycoprotein is shown in green. Candidate proteins included BGN, COL6A6, POSTN, HSPG2, LAMA4, and LAMC1; COL1 was included as a structural ECM control. The same tissue region is shown to facilitate side-by-side comparison of the spatial distributions of WFA and the candidate proteins. Pixel-based quantitative measures of protein–WFA spatial correspondence within the same region of interest are summarized in Table S8. Scale bars, 200 μm.

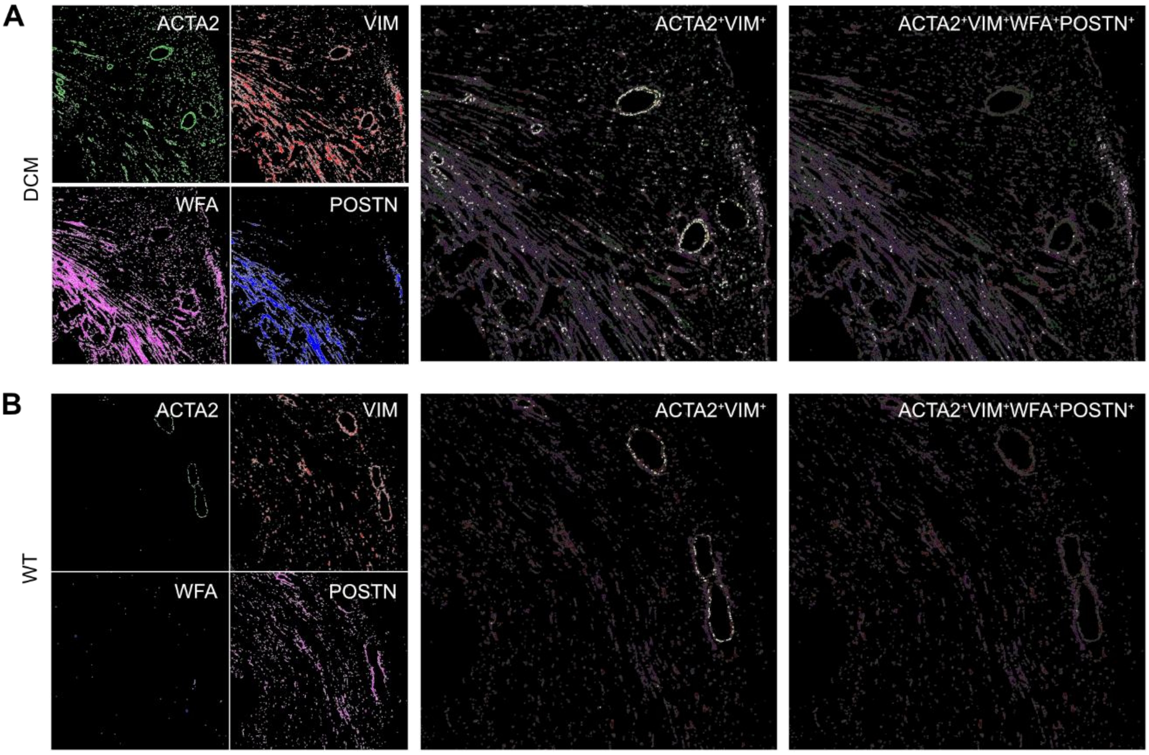


**Figure S4. HALO object colocalization maps of ACTA2, VIM, WFA, and POSTN in DCM and WT hearts.** Representative HALO outputs from (A) DCM and (B) WT heart sections are shown. Object maps for ACTA2, VIM, POSTN, and WFA are displayed together with a composite colocalization view and phenotype maps for ACTA2$^+$VIM$^+$WFA$^+$POSTN$^+$ and ACTA2$^+$VIM$^+$ objects. ACTA2$^+$VIM$^+$ objects showed a broader distribution, whereas ACTA2$^+$VIM$^+$WFA$^+$POSTN$^+$ objects were observed predominantly within WFA-enriched fibrotic regions in the representative DCM section and were limited in the WT section. Object maps were generated using the HALO Object Colocalization FL module. Detailed HALO analysis parameters and phenotype definitions are provided in Table S6.
